# Dextran sulfate sodium salt corrupted colonic crypts declined the smooth muscle tension in mouse large intestine

**DOI:** 10.1101/2020.12.17.423322

**Authors:** Sun Yiwei, Hu Aihua, Fan shouyan, Wei Lusi, Shi Yuechuan, Wen Lu, Cham Mohamed Aden, Gao Lingfeng, Wang Yang

## Abstract

Ulcerative colitis is one kind of colonic mucosa damage, shows high number of inflammatory epithelial cells. Dextran sulfate sodium salt (DSS) induce a milder onset of colitis or a more aggressive response. It may damage the protective effects on intestinal barrier. In this study, we investigated the damaging of colon crypts, evaluated the smooth muscle tension beneath corrupted crypts in DSS exposed mice.

**Methods:** female specific-pathogen-free *BALB/C* mice (n=16) are randomly divided as: group A: control mice (n=4); group B: DSS-mice (colitis, 5% DSS in drink water, days 1 to 7, n = 12). The DSS is replaced every 2 days. On day 8, mice colons are excised from the colon-cecal junction to the anus. The distal colon segment is longitude incision and aberrant crypt area are determined by methylene blue staining method. The smooth muscle strip is separated and prepared for passive tension tests. The rest segment is fixed with 10% formalin and embedded in paraffin. Histological scores are evaluated in hematoxylin-eosin staining section: crypt damage (none = 0, basal 1/3 damaged = 1, basal 2/3 damaged = 2, only the surface epithelium is intact = 3, and entire crypt and epithelium are lost = 4). The smooth muscle passive tension beneath the aberrant crypt area in DSS-mice are tested and compared with the preparations from control mice.

**Results:** In DSS uptake mice, the inflammation in large intestine mucosa damaged crypts with architectural distortions on day 7 (n=7). In crypts damage area, the smooth muscle passive tension and relative myogenic spontaneous contraction parameters are significantly reduced under the high preload conditions. The maximum rate of change of velocity of spontaneous contraction was noticeable attenuated.

**Conclusion:** Our findings demonstrate that low dosage DSS water drink result in corrupted colonic crypts. The corrupted crypts damage the large intestinal epithelium barrier, affect the smooth muscle functions, which declined in myogenic spontaneous contraction under the preload. This further may reduce the peristalsis in large intestine.

## Introduction

Dextran sulfate sodium salt (DDS) is polymer of sulfated polysaccharide, exerts chemical injury to the intestinal epithelium, resulting in exposure of the lamina propria (LP) and submucosal compartment to luminal antigens and enteric bacteria, triggering inflammation^[1, 2]^. In ulcerative colitis, denervated smooth muscle tends to show an alteration in muscle tension responsiveness to the stimuli that due to abnormal mobilization of intracellular calcium. The identification of this abnormality may provide a potential avenue for future understanding of ulcerative colitis. In ulcerative colitis, crypt dropout, basal plasmacytosis with lymphoid aggregates, and giant cells in the lower region of the lamina propria was the typical morphological changes, which was accompanied by extension of submucosal smooth muscle bands between glands. Lamina propria forms the connective tissue core surrounding the crypt epithelium. The crypt and the lamina propria are separated by a distinct basement membrane composed of an ultrastructural apparent basal lamina and a deeper network of collagenous fibers^[3]^. The pericryptal sheath surrounding colonic crypts is an effective barrier both to dextran movement. There is a greater accumulation of dextran in the crypt lumens of descending colon than in the caecal crypts whereas no such structure surrounds the caecal crypts^[4]^. Low dietary Na¤ intake raised rat plasma aldosterone and stimulated distal pericryptal sheath growth and adhesiveness as shown by increased amounts of F_actin, smooth muscle actin, ß-catenin and E-cadherins in the pericryptal zone^[5]^. The accumulation of smooth muscle actin was relative to the muscle spasms in persistent in involuntary muscle contractions^[6]^, ß-catenin is reported function as a novel mediator of glucose transport in muscle and may contribute to insulin-induced actin-cytoskeleton remodelling to support GLUT4 translocation^[7]^, which exerts a major effect on smooth muscle contractile and relaxation responses^[8]^. The increasing of smooth muscle tone, that arises from disrupted crypt architecture, was thought through the distinct mechanotransductive signaling mechanisms^[9]^. The determination of maximum velocity of shortening was depending on the mechanical preload on the muscle^[10]^. Muscle’s passive tension arises from elastic spring-like elements stretched beyond their resting length. The characteristic of smooth muscles is that at the length which will give the maximal active tension, they characteristically have considerable passive tension. This is, aside from striation, the big difference between smooth and skeletal muscle.

The large intestine circular smooth muscle layer causes its wall to form haustra which disappeared when muscle tone was lost. Its contraction causes the food to be churned and maximizing absorption. The smooth muscle passive tension (PT), which is the passive stiffness component, is the mechanical compound to initiate the circular smooth muscle contraction. The large intestine smooth muscle are regulated by biochemical pathways and represents intracellular crosslinks. Foci of lamina propria inflammation, edema, aphthous ulcers, and focal crypt injury produce an irregular distribution of crypts in the lamina propria. This may infect the smooth muscle, affect the activity of smooth muscle. A decrease in the intracellular concentration of activator Ca^2+^ elicits smooth muscle cell relaxation. Several mechanisms are implicated in the removal of cytosolic Ca^2+^ and involve the sarcoplasmic reticulum and the plasma membrane. Ca^2+^ uptake into the sarcoplasmic reticulum is dependent on ATP hydrolysis. The aberrant crypt altered calcium signaling in colonic smooth muscle^[11]^.

In this study we investigated DDS water uptake induced aberrant crypt in mice, evaluated the smooth muscle passive tension and relative myogenic spontaneous contraction beneath the aberrant crypt.

## Methods

### Animal

A total of 16 specific-pathogen-free *BALB/C* mice (aged 8 to 10 weeks, weighing 20–24g) were purchased from the Laboratory Animal Center of Hainan Medical College (Hainan island, China), maintained in clean cages under a 12h light-dark cycle and conventional housing conditions, fed with standard mouse chow. All animal experiments were performed in accordance with the National Institutes of Health Guide for the Care and Use of Laboratory Animals, and the protocol was approved by the Animal Ethics Committee of Hainan Medical College (Approval ID: Q20170013). 5% DSS (0216011080, MW 36–50 kDa, MP Biomedicals) in drinking water was used to induce acute colitis. The DSS was replaced every 2 days. Mice were randomly divided and treated as follows: group A (normal, n = 4): mice received sham (saline, days 1 to 14); group B (DSS uptake, n = 12). At day 15, all animals were euthanatized.

### The determination of large intestine aberrant crypt area

The large intestine was excised from the colon-cecal junction to the anus, and the lengths of the colon were measured. After the mice large intestines were removed from the abdominal cavity. The segment of the distal segment was taken and immediately immersed in *Ringer’s* solution (pH7.4). The segment was transverse axis midline incision. The internal luminal wall and the surface of the villus epithelium was stained by methylene blue staining method. The aberrant crypt area referenced the method of Gupta AK^[12]^, McGinley JN^[13]^. After staining, the aberrant crypt area was easily distinguished from surrounding normal mucosa in intact large intestine wall under the microscope. The large intestine wall of aberrant crypt area was obtained by ophthalmic scissors and prepared for the further analysis.

### The histological scores

The distal segment was fixed with 10% formalin and embedded in paraffin. Paraffin sections (4 μm) were stained with hematoxylin-eosin (H&E). Histological scores were evaluated as follow: inflammation (none=0, slight=1, moderate=2, and severe=3), inflamed area/extent (mucosa=1, mucosa and submucosa=2, and transmural=3), crypt damage (none=0, basal 1/3 damaged=1, basal 2/3 damaged=2, only the surface epithelium is intact=3, and entire crypt and epithelium are lost=4), and percent involvement (1–25% = 1, 26–50% = 2, 51– 75% = 3, and 76–100% = 4).

### The large intestine smooth muscle tissue preparation

The smooth muscle layer of aberrant crypt area was separated from the entire colonic mucosa in large intestine wall using Adson forceps. The smooth muscle strip endings were ligatured and stable in *Ringer’s* solution for further analysis.

### The smooth muscle strip passive tension measurement

The passive tension and myogenic spontaneous contraction of longitude smooth muscle strips beneath aberrant crypt, and non-aberrant crypt area were evaluated. The strips were one end fixed on the tension transducer (model JZ-100, Beijing institute of aerospace medical engineering, Beijing, China), the other end was fixed on micro step tuning. The transducer was connected to the physiological polygraph device (BL-420S, Chengdu Taimeng software Co. Ltd., China). The strips were longitude stretched to obtain a 1gram preload (1g, low preload), then rapid stretch to obtain a passive tension curve. The muscle strip was further slowly longitude stretched to obtain 5gram (5g, high preload), then rapid stretch to obtain the passive tension curve under the high preload. Under the preload condition, in each rapid stretch the muscle strip was extended 0.1mm on its longitude axis, and stretched 5 time continually. The passive tension and relaxation period after each rapid stretch were recorded. The myogenic spontaneous contraction amplitude (A_s_), maximum contraction velocity (V_max_), maximum instantaneous contraction force (F_max_) were analyzed. The data were compared with control mice.

## Statistics

The large intestine crypts images are representative of at least three independent experiments. The mean value of myogenic spontaneous contraction A_s_, V_max_, F_max_ from 5 steps and standard error of mean (SEM) was calculated and compared between DSS uptake and control mice. P values were calculated by Mann-Whitney U Test (Microsoft Excel 2019, version 1808). P value < 0. 05 was considered significant.

## Results

### The histological changes

**Figure 1a** is the large intestine distal segment colitis sample from the DSS-mice. In control mice, the large intestine consists of a crypt/villus and crypt/surface epithelium unit, respectively. The bulk of the villus and surface epithelium is composed of differentiated columnar epithelial cells that are divided into absorptive cells for enterocytes and secretory cells. The locations of the multi-potent stem cells where the crypt progenitor cells differentiated to epithelial cells are located in the crypt compartment (**Figure 1b**, red Square frame). In DSS-mice, the crypt image features showed significant differences to the control mice. **Figure 1c** shows examples of histology of DSS-mice aberrant crypt. The crypt structures tend to be not uniform in size and loss the general shape across the field of view. The crypt tubular shape in transverse view is lost the regular. Collagen distribution appears relatively even throughout the field of view. The crypt bottom has early dysplasia, and tend to vary in size across the field of view.

**Fig. 1.**
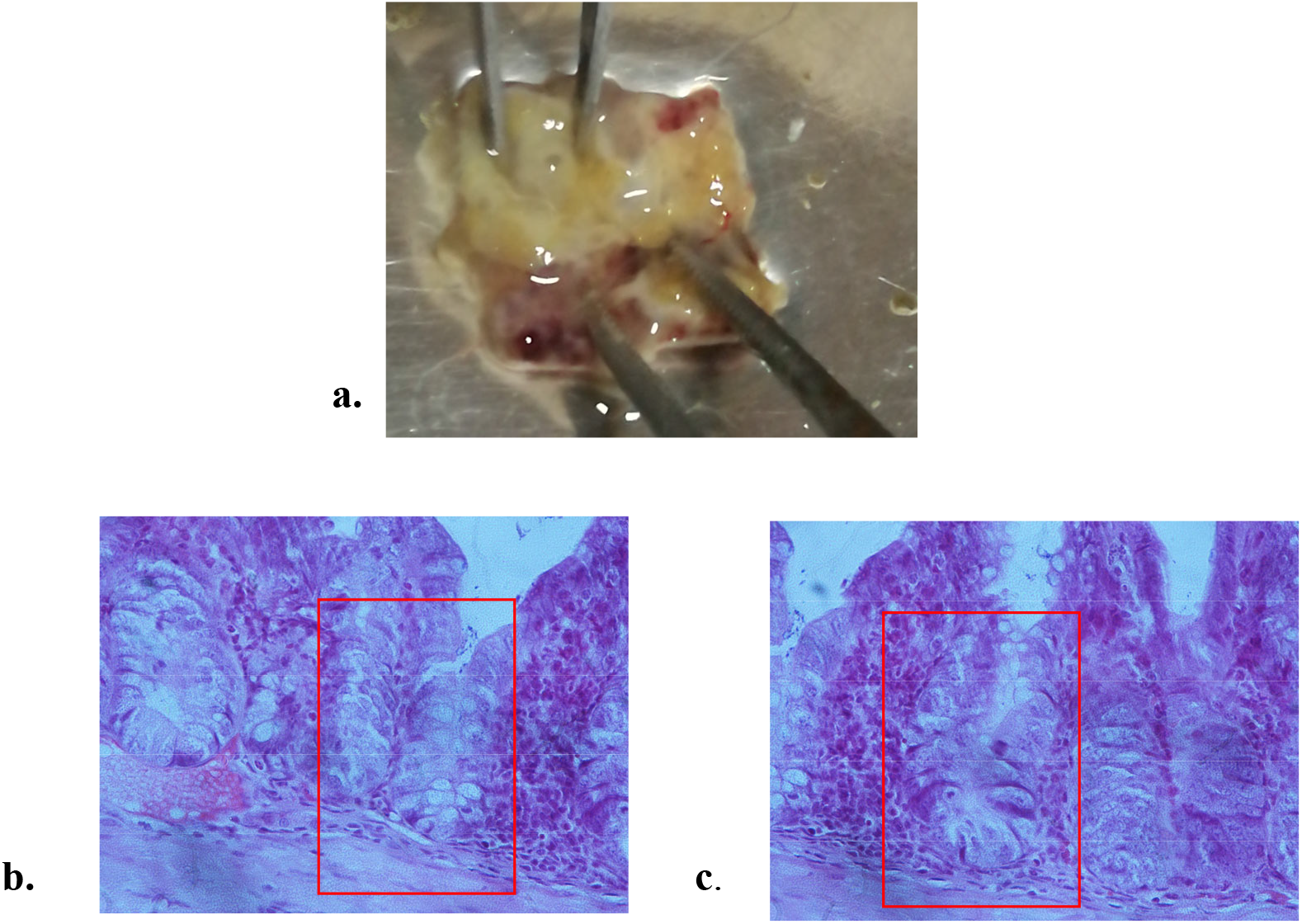
The colitis and crypts changes of the large intestinal mucosa in DSS-mice

The histological scores in DSS-mice distal segment are evaluated as follow: 1) The inflammation (none, n=0; slight, n=5; moderate, n=6; severe, n=3). 2)The inflamed area or extent (mucosa, n=4; mucosa and submucosa, n=6, transmural, n=6).3) The crypt damage (none, n=2; basal damaged, n=8; only the surface epithelium is intact, n=3; entire crypt and epithelium are lost, n=1). 4) The percent involvement (1–25%, n=7, 26–50%, n=5, 51– 75%, n=2, 76–100%, n=0). The results are summarized in **Table 1**.

**Table 1.** The large intestine wall histological score in DSS uptake mice (**n=12**)

| Inflammation |  | Inflamed area/extent |  | Crypt damage |  | Percent involvement |  |
| --- | --- | --- | --- | --- | --- | --- | --- |
| none | 0 | mucosa | 4 | none | 2 | 1–25% | 7 |
| slight | 5 | mucosa/submucosa | 6 | basal 1/3 damaged | 4 | 26–50% | 5 |
| moderate | 6 |  |  | basal 2/3 damaged | 4 | 51– 75% | 2 |
|  |  |  |  | Only surface epithelium intact | 3 | 76–100% | 0 |
| severe | 3 | transmural | 6 | entire crypt/epithelium lost | 1 |  |  |

### The smooth muscle spontaneous contraction beneath aberrant crypt area

The toxic colitis occurs when inflammation extends into the smooth muscle layer of the intestinal wall. In the smooth muscle preparation that obtained from the aberrant crypt area, we tested the rapid stretch induced myogenic spontaneous contraction under the preload conditions.

When the muscle strip lengthens to obtain a 1gram (1g, initial preload), the initial passive tension curve has no difference between DSS treated and control preparation. The significant difference is observed after rapid stretch and the smooth muscle bearing the lengthening after the stretch. In control mice, under the low preload condition, each rapid stretch induce a myogenic spontaneous contraction that followed the maximum passive tension (**Figure. 2a**), however, the amplitude of myogenic spontaneous contraction have variation after each stretch (the * marked peak wave). Under the high preload condition, each rapid stretch induce significant myogenic spontaneous contraction that tightly closed to the maximum passive tension peak wave (**Figure. 2b**), however, the amplitude of myogenic spontaneous contraction have significantly incerased after each stretch (the * marked peak wave).

**Fig. 2.**
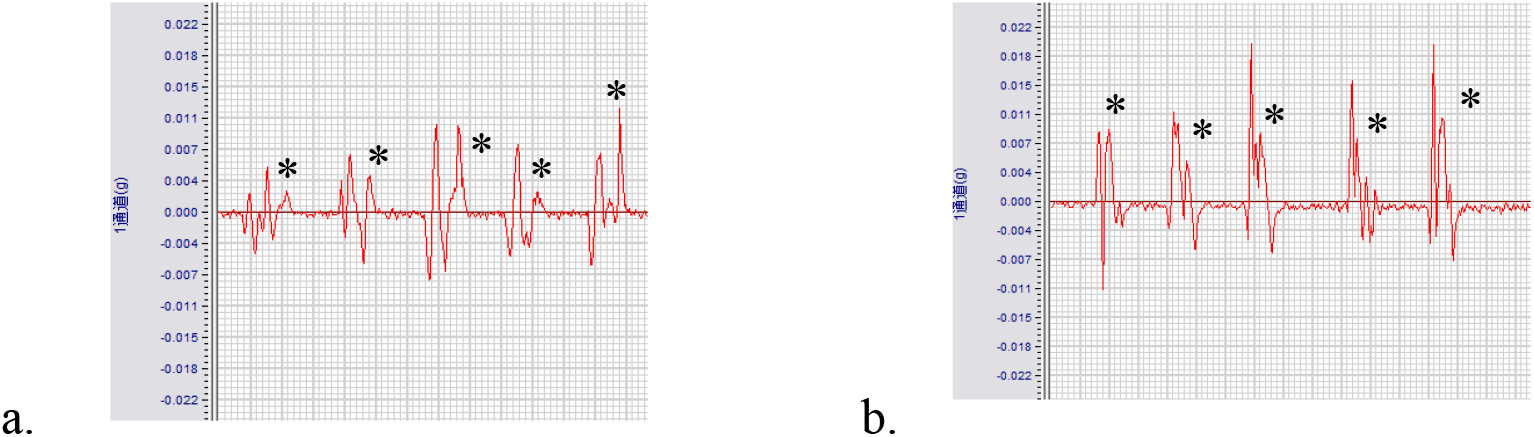
The rapid stretch induced myogenic spontaneous contraction

Under the low preload, the rapid stretching induced a significant passive tension (PT_max_) in control mice, however the PT_max_ is significantly reduced in DSS-mice. The PT_max_ ratio between high preload and low preload is significantly reduced (**Figure. 3a**). The myogenic spontaneous contraction amplitude As ratio between high preload and low preload is significantly low in DSS-mice (**Figure. 3b**).

**Fig 3.**
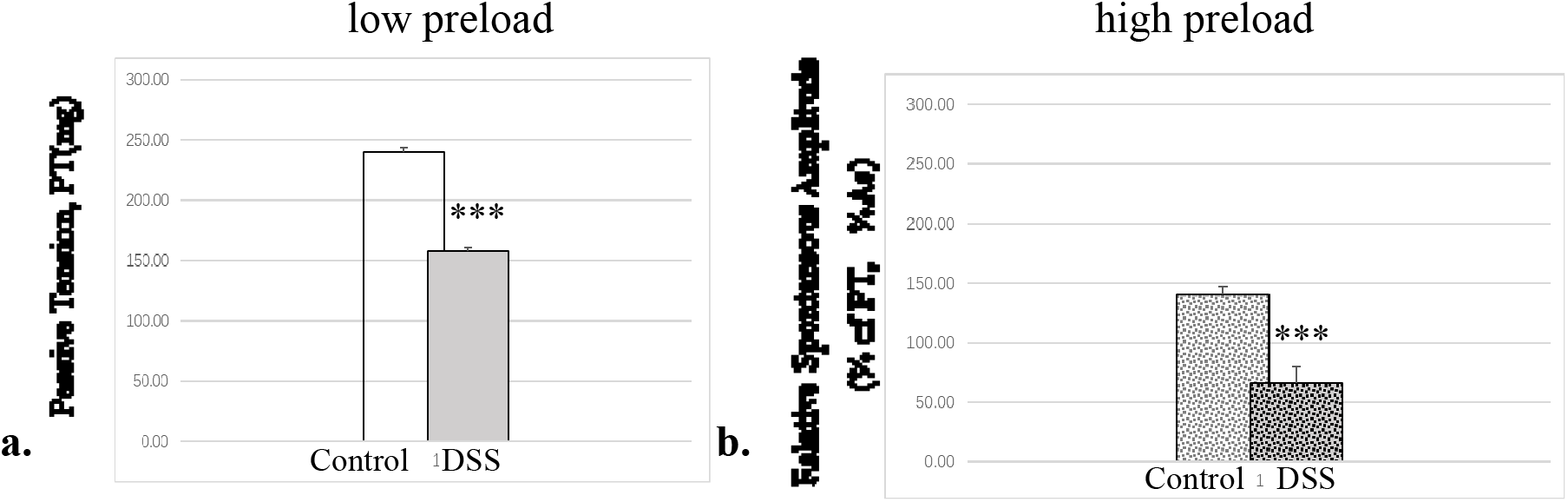
The smooth muscle passive tension amplitude in DSS-mice

The rapid stretch induced maximum velocity of myogenic spontaneous contraction (V_max_) is significantly low in the high preload preparation (compare with the control mice, **Figure 4b**, the dot gray column). However, the maximum velocity of myogenic spontaneous contraction (V_max_) have no significantly changes in the low preload preparation (compare with the control mice, **Figure 4a**, the gray column). Meanwhile, the maximum instantaneous contraction force (F_max_) of the myogenic spontaneous contraction have significantly reduced under the high preload condition (**Figure 4d**, the dot gray column),. However, the F_max_ of myogenic spontaneous contraction have no significantly changes in the low preload preparation (compare with the control mice, **Figure 4c**, the gray column). The V_max_ and F_max_ of control and DSS-mice are 38.99±6.95 to 38.39±4.82 g/sec (***, p<0.001) and 101.78±67.41 to 62.50±12.31 g/sec (***, p<0.001), respectively.

**Fig. 4.**
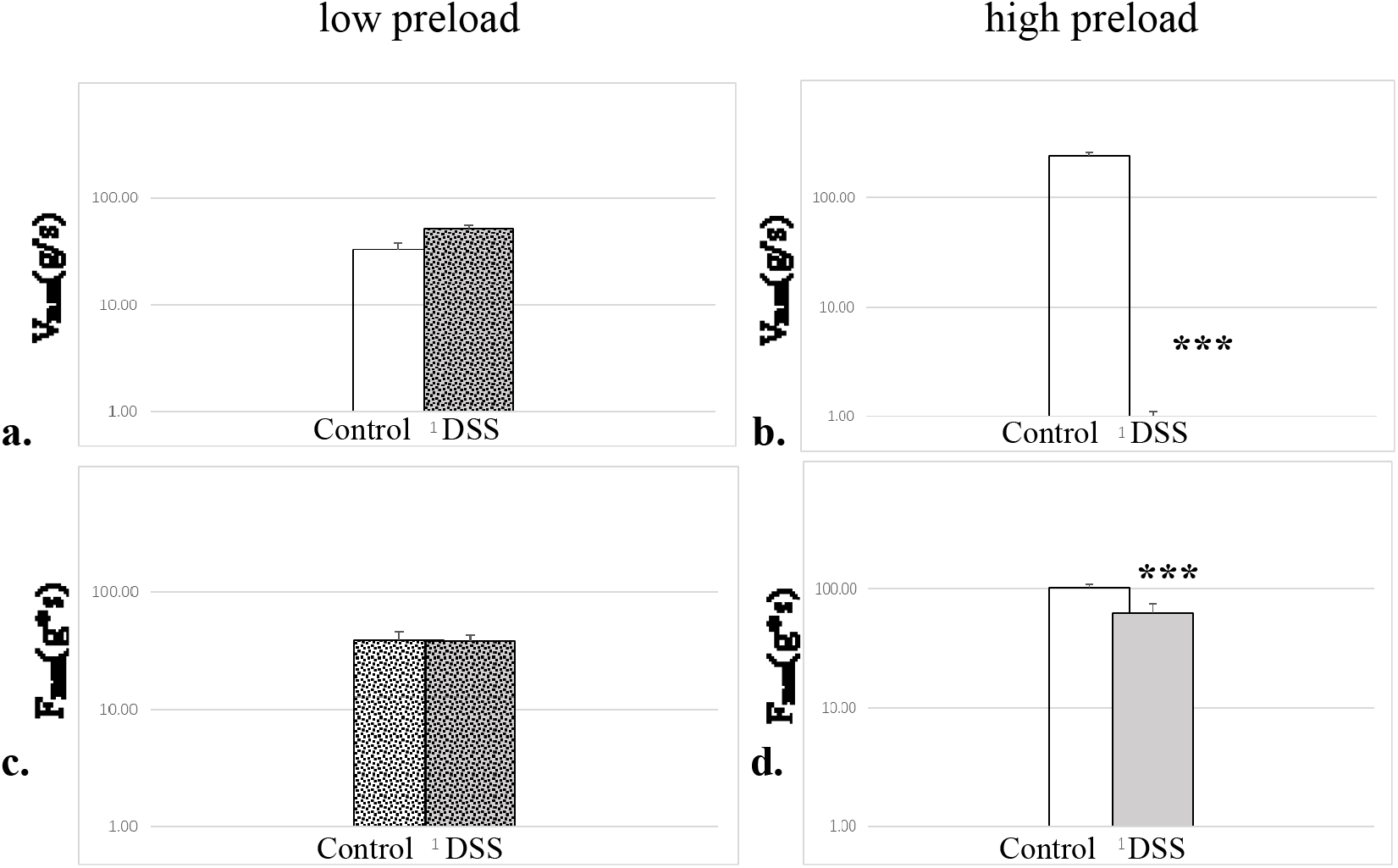
The myogenic spontaneous contraction in DSS-mice

The V_max_ and F_max_ significantly reduced in DSS-mice indicated that smooth muscle myogenic spontaneous contraction (active contraction) was weakening beneath the aberrant crypt area.

## Discussion

The intestinal epithelium withstands continuous mechanical, chemical and biological insults despite its single-layered, simple epithelial structure. The crypt-villus architecture in combination with rapid cell turnover enables the intestine to act both as a barrier and as the primary site of nutrient uptake. Constant tissue replenishment is fueled by continuously dividing stem cells that reside at the bottom of crypts. Dextran sulphate sodium (DSS) induced imbalance in key signaling pathways can cause the initiation of large intestine disease. DSS induced colitis is a reproducible model that morphologically and symptomatically resembles ulcerative colitis in human. This compound mainly affects the large intestine, especially middle and distal third of large intestine^[14]^. Most of the reports suggested that DSS causes erosions with complete loss of surface epithelium because of its direct toxic effect on epithelial cells. In the mammalian intestine, crypts of Leiberkühn house intestinal epithelial stem/progenitor cells at their base. During homeostasis, differentiated colonocytes metabolized butyrate likely preventing colitis from reaching proliferating epithelial stem/progenitor cells within the crypt. Exposure of stem/progenitor cells *in vivo* to butyrate through either mucosal injury or application to a naturally crypt-less host organism led to inhibition of proliferation and delayed wound repair^[15]^.

DSS dosage is a key factor in mucosa damaging. 1% DSS for 9 days and 2% DSS for 6 days minimum induces colitis in male wild type rats^[16]^. The severity of colitis i.e. mild, moderate, severe colitis may be varied by varying the DSS treatment period. Administering 3%, w/v DSS for 7 days and sacrificing on the 8th day induces mild colitis whereas moderate colitis is induced by administering 3% DSS for a period of 14 days i.e. from days 1 to 7 and 22 to 28 and then sacrificing animals on the 29th day^[17]^. Our results suggested that the low dosage of DSS uptake through the drink water damaged crypts. The histological score indicated that more than half of the DSS-mice have the morphological damage in distal segment (n=8), 12 of the DSS-mice were submucosa transmural inflamed. This perhaps induced by reducing in *Lactobacillus sp*. and protective short-chain fatty acid production, and alter gut immune homeostasis and lead to increased vulnerability to inflammatory insults^[18]^.

The toxic colitis occurs when inflammation extends into the smooth muscle layer of the intestinal wall, paralyzing the colon muscle. This may lead to colon dilatation, and sometimes perforation. In physiological intestine, the frequency of contractions was controlled by constitutive nitric oxide. In colitis induced transient increases in the amplitude of spontaneous contractions coincident with a loss of nitric oxide synthase activity. The initial colitis induced a remodeling of the neural control of spontaneous contractions reflecting changes in their regulation by constitutive nitric oxide synthase and iNOS^[19]^. In this study the denervation smooth muscle is prepared for investigating the myogenic contraction. The smooth muscle layer beneath the aberrant crypt area are significantly reduction of its passive tension under the high preload condition. This indicated the low stiffness of the smooth muscle after DSS toxic inflammation and the aberrant crypt formation. This evidence may relative to the colon dilatation and perforation in the toxic colitis.

The smooth muscle myogenic contraction can be demonstrated in the autoregulation of the cavity organs. The studies of urinary bladder detrusor indicate that spontaneous contraction is responsible for regenerating adjustable preload stiffness^[20]^ and for length adaptation^[21]^. Strength of the myogenic response is greatest when the intestine wall opposite to the intraluminal pressure and the enlargement of the lumen. Strength varies in different intestine segment. The different of the myogenic response also observed in variation of internal environment. In this study, passive tension amplitude is responsible for the preload conditions, and the myogenic spontaneous contraction is responsible for the smooth muscle beneath the aberrant crypt area. Both in control mice and DSS-mice, the large intestine smooth muscle exhibits the phasic myogenic contraction independent of neural input. In addition, the amplitudes of contraction are muscle length dependent, and amplitude increasing at longer muscle lengths. The myogenic retrogression response is consistent with the DSS toxicity induced crypt damaging.

## Acknowledgement

This study was sponsored by Hainan provincial college student innovation and entrepreneurship project (No. S201911810023).

## Competing Interests

No potential competing interests.

## Notes

### Competing Interest Statement

The authors have declared no competing interest.

